# A direct approach to estimating false discovery rates conditional on covariates

**DOI:** 10.1101/035675

**Authors:** Simina M. Boca, Jeffrey T. Leek

## Abstract

Modern scientific studies from many diverse areas of research abound with multiple hypothesis testing concerns. The false discovery rate is one of the most commonly used error rates for measuring and controlling rates of false discoveries when performing multiple tests. Adaptive false discovery rates rely on an estimate of the proportion of null hypotheses among all the hypotheses being tested. This proportion is typically estimated once for each collection of hypotheses. Here we propose a regression framework to estimate the proportion of null hypotheses conditional on observed covariates. This may then be used as a multiplication factor with the Benjamini-Hochberg adjusted p-values, leading to a plug-in false discovery rate estimator. Our case study concerns a genome-wise association meta-analysis which considers associations with body mass index. In our framework, we are able to use the sample sizes for the individual genomic loci and the minor allele frequencies as covariates. We further evaluate our approach via a number of simulation scenarios.

## 1 Introduction

Multiple testing is a ubiquitous issue in modern scientific studies. Microarrays (Brown, 1995), next-generation sequencing (Shendure and Ji, 2008), and high-throughput metabolomics (Lindon et al., 2011) make it possible to simultaneously test the relationship between hundreds or thousands of biomarkers and an exposure or outcome of interest. These problems have a common structure consisting of a collection of variables, or features, for which measurements are obtained on multiple samples, with a hypothesis test being performed for each feature.

When performing thousands of hypothesis tests, the most widely used framework for controlling for multiple testing is the false discovery rate (FDR). For a fixed unknown parameter *μ*, and testing a single null hypothesis *H*_0_: *μ* = *μ*_0_ versus some alternative hypothesis, for example, *H*_1_: *μ* = *μ*_1_, the null hypothesis may either truly hold or not for each feature. Additionally, the test may lead to *H*_0_ either being rejected or not being rejected. Thus, when performing *m* hypothesis tests for m different unknown parameters, Table 1 shows the total number of outcomes of each type, using the notation from Benjamini and Hochberg (1995). We note that *U*, *T*, *V*, and *S*, and as a result, also *R* = *V*+ *S*, are random variables, while *m*_0_, the number of null hypotheses, is fixed and unknown.

**Table 1.** Outcomes of testing multiple hypotheses.

|  | Fail to reject null | Reject null | Total |
| --- | --- | --- | --- |
| Null true | $U$ | $V$ | $m_0$ |
| Null false | $T$ | $S$ | $m - m_0$ |
| | $m - R$ | $R$ | $m$ |

The FDR, introduced in Benjamini and Hochberg (1995), is the expected fraction of false discoveries among all discoveries. The false discovery rate depends on the overall fraction of null hypotheses, namely 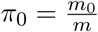. This proportion can also be interpreted as the *a priori* probability that a null hypothesis is true, *π*_0_.

When estimating the FDR, incorporating an estimate of *π*_0_ can result in a more powerful procedure compared to the original Benjamini and Hochberg (1995) procedure (BH); moreover, as *m* increases, the estimate of *π*_0_ improves, which means that the power of the multiple-testing approach does not necessarily decrease when more hypotheses are considered (Storey, 2002).

Most modern adaptive false discovery rate procedures rely on an estimate of *π*_0_ using the data of all tests being performed. But additional information, in the form of meta-data, may be available to aid the decision about whether to reject the null hypothesis for a particular feature. We focus on an example from a genome-wide association study (GWAS) meta-analysis, in which millions of genetic loci are tested for associations with an outcome of interest - in our case body mass index (BMI) (Locke et al., 2015). Different loci may not all be genotyped in the same individuals, leading to loci-specific sample sizes. Additionally, each locus will have a different population-level frequency. Thus, the sample sizes and the frequencies may be considered as covariates of interest. Other examples exist in set-level inference, including gene-set analysis, where each set has a different fraction of false discoveries. Adjusting for covariates independent of the data conditional on the truth of the null hypothesis has also been shown to improve power in RNA-seq, eQTL, and proteomics studies (Ignatiadis et al., 2016).

In this paper, we seek to better understand the impact of sample sizes and allele frequencies in the BMI GWAS data analysis by building on the approaches of Benjamini and Hochberg (1995), Efron et al. (2001), and Storey (2002) and the more recent work of Scott et al. (2015), which frames the concept of FDR *regression* and extends the concepts of FDR and *π*_0_ to incorporate covariates, represented by additional meta-data. Our focus will be on estimating the covariate-specific *π*_0_, which will then be used as a plug-in estimator when estimating the false discovery rate, similar to the work of Storey (2002). We will also show how this can be seen as an extension of our work (Boca et al., 2013) on set-level inference, where an approach which focused on estimating the fraction of non-null variables in a set was developed, introducing the idea of “atoms” - non-overlapping sets based on the original annotations - and the concept of the “atomic FDR.” We thus provide a more direct approach to estimating the FDR conditional on covariates and compare our estimates to those of Scott et al. (2015), as well as to the BH and Storey (2002) approaches.

In Section 2 we introduce the motivating case study, a BMI GWAS meta-analysis, which will be discussed throughout the paper. In Section 3, we review the definitions of FDR and *π*_0_ and their extensions to consider conditioning on specific covariates. In Section 4, we discuss estimation and inference procedures in our FDR regression framework and apply them to the GWAS case study. In Section 5, we consider special cases within this framework, including how the no covariates case and the case where the features are partitioned are related to the “standard” estimation procedures. In Section 6, we explore some theoretical properties of the estimator, including showing that, under certain conditions, it is an asymptotically conservative estimator of the covariate-level *π*_0_. In Section 7 we consider results from a variety of simulation scenarios. Finally, Section 8 provides our statement of reproducibility and Section 9 provides the discussion.

## 2 Motivating case study: adjusting for sample size and allele frequency in GWAS meta-analysis

As we have described, there are a variety of situations where meta-data could be valuable for improving the decision of whether a hypothesis should be rejected in a multiple testing framework, our focus being on an example from the meta-analysis of data from GWAS for BMI (Locke et al., 2015). Using standard approaches such as Storey (2002) we can estimate the fraction of single nucleotide polymorphisms (SNPs) - genomic positions (loci) which show between-individual variability - which are not truly associated with BMI and use it in an adaptive FDR procedure. However, our proposed approach allows further modeling of this fraction as a function of additional study-level meta-data.

In a GWAS, data are collected for a large number of SNPs in order to assess their associations with an outcome or trait of interest (Hirschhorn and Daly, 2005). Each person usually has one copy of the DNA at each SNP inherited from their mother and one from their father. At each locus there are usually one of two types of DNA, called alleles, that can be inherited, which we denote *A* and *a*. In general, *A* refers to the variant that is more common in the population being studied and *a* to the variant that is less common, usually called the minor allele. Each person has a genotype for that SNP of the form *AA*, *Aa*, or *aa*. For example, for a particular SNP, of the 4 possible DNA nucleotides, adenine, guanine, cytosine, and thymine, an individual may have either a cytosine (C) or a thymine (T) at a particular locus, leading to the possible genotypes CC, CT, and TT. If the C allele is less common in the population, then C is the minor allele. The number of copies of *a*, which is between 0 and 2, - is often assumed to follow a binomial distribution, which generally differs between SNPs.

Typically, a GWAS involves performing an association test between each SNP and the outcome of interest by using a regression model, including the calculation of a p-value. While GWAS studies are often very large, having sample sizes of tens of thousands of individuals genotyped at hundreds of thousands of SNPs, due to the small effect sizes being detected, meta-analyses combining multiple studies are often considered (Neale et al., 2010; Hirschhorn and Daly, 2005). In these studies, the sample size may not be the same for each SNP, for example if different individuals are measured with different technologies which measure different SNPs. Sample size is thus a covariate of interest, as is the minor allele frequency (MAF) of the population being studied, which will also vary between SNPs. The power to detect associations increases with MAF. This is related to the idea that logistic regression is more powerful for outcomes that occur with a frequency close to 0.5. Our approach will allow us to better quantify this dependence in order to guide the planning of future studies and improve understanding of already-collected data.

We consider data from the Genetic Investigation of ANthropometric Traits (GIANT) consortium, specifically the genome-wide association study for BMI (Locke et al., 2015). The GIANT consortium performed a meta-analysis of 339,224 individuals measuring 2,555,510 SNPs and tested each for association with BMI. 322,154 of the individuals considered in Locke et al. (2015) are of European descent and the study uses the HapMap CEU population - which consists of individuals from Utah of Northern and Western European ancestry (Frazer et al., 2007) - as a reference. We used the set of results from the GIANT portal at http://portals.broadinstitute.org/collaboration/giant/index.php/GIANT_consortium_data_files, which provides the SNP names and alleles, effect allele frequencies (EAFs) in the HapMap CEU population and results from the regression-based association analyses for BMI, presented as beta coefficients, standard errors, p-values, and sample size for each SNP.

We removed the SNPs that had missing EAFs, leading to 2,500,573 SNPs. For these SNPs, the minimum sample size considered was 50,002, the maximum sample size 339,224, and the median sample size 235,717 - a relatively wide range. Figure 1 shows the dependence of p-values on sample sizes within this dataset. As we considered the MAF to be a more intuitive covariate than the effect allele frequency (EAF), we also converted EAF values > 0.5 to MAF=1-EAF and changed the sign of the beta coefficients for those SNPs. The MAFs spanned the entire possible range from 0 to 0.5, with a median value of 0.208.

**Figure 1:**
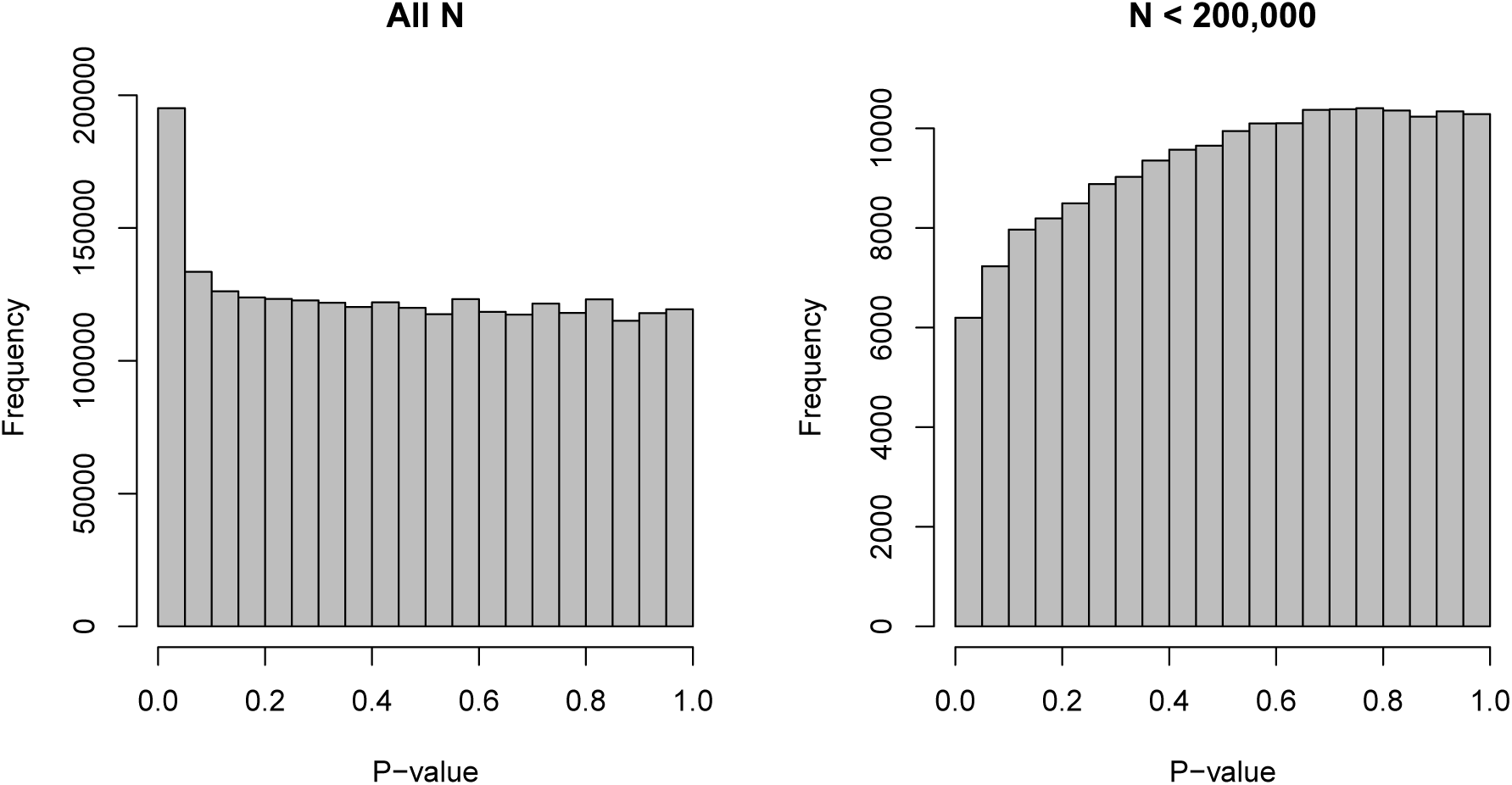
Histograms of p-values for the SNP-BMI tests of association from the GIANT consortium. Panel a) shows the distribution for all sample sizes N (2,500,573 SNPs), while panel b) shows the subset N <200,000 (187,114 SNPs).

## 3 Covariate-specific *π*_0_ and FDR

We will now review the main concepts behind the FDR and the *a priori* probability that a null hypothesis is true, and consider the extension to the covariate-specific FDR, and the covariate-specific *a priori* probability. A natural mathematical definition of the FDR would be:

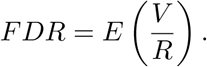

However, *R* is a random variable that can be equal to 0, so the definition that is generally used is:

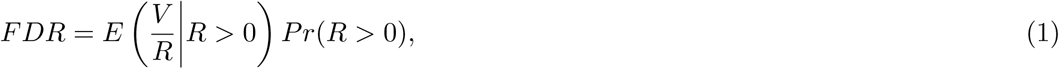

namely the expected fraction of false discoveries among all discoveries multiplied by the probability of making at least one rejection.

We index the *m* null hypotheses being considered by 1 ≤ *i* ≤ *m*: *H*_01_, *H*_02_, …, *H*_0*m*_. For each *i*, the corresponding null hypothesis *H*_0*i*_ can be considered as being about a binary parameter *θ*_*i*_, such that:

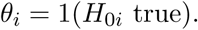

Thus, assuming that *θ*_*i*_ are identically distributed, the *a priori* probability that a feature is null is:

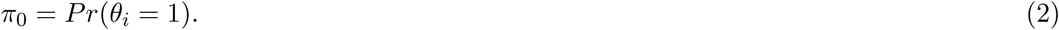

For the GWAS meta-analysis dataset, π_0_ represents the proportion of SNPs which are not truly associated with BMI or, equivalently, the prior probability that any of the SNPs is not associated with BMI.

We now extend the definitions of *π*_0_ and FDR to consider conditioning on a set of covariates concatenated in a column vector X_*i*_ of length *c*, possibly with *c* = 1:

### Definition 1

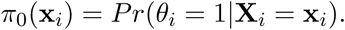

### Definition 2

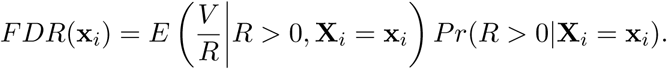

## 4 Estimation and inference for covariate-specific *π*_0_ and FDR in a regression framework

We will now discuss the estimation and inference procedures for *π*_0_(x_*i*_) and FDR(x_*i*_). We assume that a hypothesis test is performed for each *i*, summarized by a p-value *P*_*i*_. Our approach is based on thresholding the p-values at a given *λ* ∈ (0,1), resulting in binary indicators *Y*_*i*_ = 1(*P*_*i*_ > *λ*). These are then treated as outcomes in a regression model. In the rest of the section, we show how this procedure allows us to estimate *π*_0_(x_*i*_) and FDR(x_*i*_).

Since *Y*_*i*_ is a dichotomous random variable that is 1 when the null hypothesis *H*_0*i*_ is not rejected at a significance level of *λ* and 0 when it is rejected,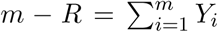 for a fixed, given *λ*. The null p-values will come from a Uniform(0,1) distribution, while the p-values for the features from the alternative distributions *G*_X*i*_, defined as:

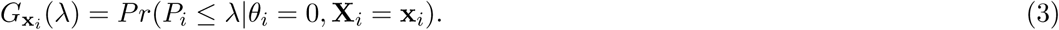

The major assumption we make moving forward is that *conditional on the null, the p-values do not depend on the covariates.* This means that the probability of a feature being from one of the two distributions depends on the covariates but the actual test statistic and p-value under the null do not depend on the covariates further. In Theorem 3, we prove the major result we will use to derive the estimator for *π*_0_(x_*i*_).

### Theorem 3

*Suppose that m hypotheses tests are performed and that conditional on the null, the p-values do not depend on the covariates. Then:*

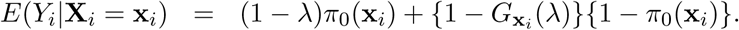

In Corollary 4, we show the corresponding result for the no-covariate case. This result is easy to prove directly, but we consider it as a corollary to Theorem 3 to show that, under the condition that *G*_X*i*_(*λ*) and *π*_0_(x_*i*_) are independent, there are no identifiability problems with the extension to covariates.

### Corollary 4

*Suppose that m hypotheses tests are performed and that conditional on the null, the p-values do not depend on the covariates and also that G*_X*i*_(*λ*) and *π*_0_(x_*i*_) *are independent. Then, assuming some distribution for the covariates:*

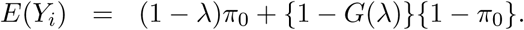

We first review the procedure which applies Corollary 4 to lead to the estimator of *π*_0_ for the no-covariate case, which is used by Storey (2002), then develop a procedure based on Theorem 3 to obtain an estimator of *π*_0_(x_*i*_). Both of them are based on assuming reasonably powered tests and a large enough *λ*, so that *G*(*λ*) ≈1 for the no-covariate case and

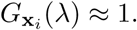

for the covariate case. Corollary 4 then leads to:

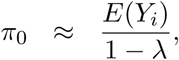

resulting in:

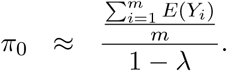

Using a method-of-moments approach, one may consider the estimator:

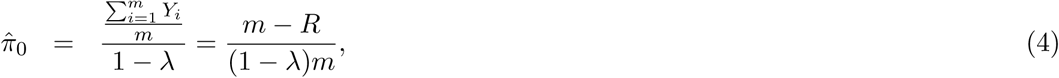

which is used by Storey (2002). For the GWAS meta-analysis dataset, using this approach with *λ* = 0.8 leads to an 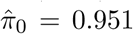 and *λ* = 0.9 to 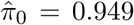. Note that in practice one may smooth over a series of thresholds, as described below; otherwise, fixed thresholds between 0.8 and 0.95 are often used. This means that *G*(*λ*) will be very close to 1, but *λ* will not be large enough to lead to numerical instability issues when dividing by 1 — *λ*.

For the covariate case, applying the same steps with Theorem 3, we get:

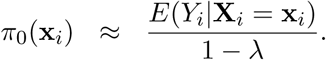

We can use a regression framework to estimate *E*(*Y*_*i*_|**X**_*i*_ = x_*i*_), then estimate *π*_0_(x) by:

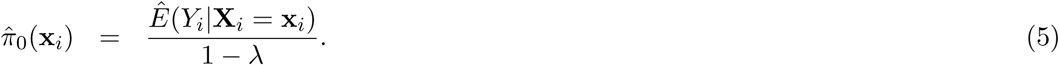

In particular, our approach will be implemented via maximum likelihood estimation of *E*(*Y*_*i*_|**X**_*i*_ = x_*i*_), assuming a logistic model. We denote by **X** the matrix of dimension *m* × (*c*+ 1), which has the *i*_*th*_ row consisting of 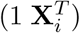. The logistic regression model will consider a *m* × *p* design matrix **Z** matrix with *p* < *m* and *rank*(**Z**) = *d* ≤ *p*, which can either be equal to **X** or include additional columns that are functions of the covariates in **X**, such as polynomial or spline terms.

A linear regression approach would be a more direct generalization of Storey (2002), but a logistic model is more natural for estimating means between 0 and 1. In particular, we note that a linear regression approach would amplify relatively small differences between large values of *π*_0_(x_*i*_), which are likely to be common in many scientific situations, especially when considering GWAS, where one may expect a relatively low number of SNPs to be truly associated with the outcome of interest.

The model we considered for the GWAS meta-analysis dataset models the SNP-specific sample size using natural cubic splines, in order to allow for sufficient flexibility. It also considers 3 discrete categories for the CEU MAFs, corresponding to cuts at the 1/3 and 2/3 quantiles, leading to the intervals [0.000,0.127) (838,070 SNPs), [0.127,0.302) (850,600 SNPs), and [0.302,0.500] (811,903 SNPs).

Note that thus far we have considered the estimate of *π*_0_(x_*i*_) at a single threshold *λ*, so that 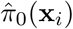 is in fact 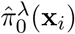. We can consider smoothing over a series of thresholds to obtain the final estimate, as done by Storey and Tibshirani (2003). In particular, in the remainder of this manuscript, we used cubic smoothing splines with 3 degrees of freedom over the series of thresholds 0.05, 0.10, 0.15,…, 0.95, following the example of the qvalue package (Storey et al., 2015), with the final estimate being the smoothed value at *λ* = 0.95. The estimates should generally be thresholded at 1, as Eq. (5) may otherwise lead to values greater than 1. It is also possible but less likely that the smoothed estimate would be below 0, hence we also threshold it at 0.

If we assume that the p-values are independent, we can also use bootstrap samples of them to obtain a confidence interval for 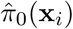. The details for the entire estimation and inference procedure for *π*_0_(x_*i*_) are in Algorithm 1.

### 4.1

#### Algorithm 1 Estimation and inference for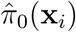

(a) Obtain the p-values *P*_1_, *P*_2_,…,*P*_*m*_, for the *m* hypothesis tests.

(b) For a given threshold *λ*, obtain *Y*_*i*_ = 1(*P*_*i*_ >*λ*) for 1 ≤*i* ≤ *m*.

(c) Estimate *E*(Y_*i*_|**X**_*i*_ = x_*i*_) via logistic regression using a design matrix **Z** and *π*_0_(x_*i*_) by:

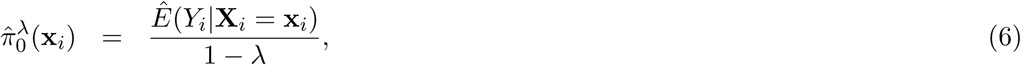

thresholded at 1 if necessary.

(d) Smooth 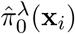 over a series of thresholds *λ* ∈ (0,1) to obtain 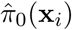, by taking the smoothed value at the largest threshold considered. Take the minimum between each value and 1 and the maximum between each value and 0.

(e) Take *B* bootstrap samples of *P*_1_, *P*_2_,…, *P*_*m*_ and calculate the bootstrap estimates 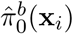 for 1 ≤ *b* ≤ *B* using the procedure described above.

(f) Form a 1 — *α* confidence interval for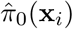 by taking the 1 — *α*/2 quantile of the 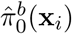 as the upper confidence bound, the lower confidence bound being *α*/2.

In order to estimate FDR(x_*i*_), we multiply the BH adjusted p-values by 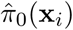, thus leading to a simple plug-in estimator, denoted 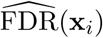. This is done in the spirit of Storey (2002), whose approach uses an estimate which is not conditional on covariates.

Figure 2 shows the estimates of *π*_0_(x_*i*_) plotted against the SNP-specific sample size N for the data analysis, stratified by the CEU MAFs for a random subset of 50,000 SNPs. We note that the results are similar for *λ* = 0.8, *λ* = 0.9, and for the final smoothed estimate. A 95% bootstrap confidence interval based on 100 iterations is also shown for the final smoothed estimate. Our approach is compared to that of Scott et al. (2015), which assumes that the test statistics are normally distributed. We considered both the theoretical and empirical null Empirical Bayes (EB) estimates of Scott et al. (2015), implemented in the `FDRreg` package (Scott et al., 2015). The former assumes a N(0,1) distribution under the null, while the latter estimates the parameters of the null distribution. Both approaches show similar qualitative trends to our estimates, although the empirical null tends to result in much higher values over the entire range of N, while the theoretical null leads to lower values for smaller N and larger or comparable values for larger N. Our results are consistent with intuition - larger sample sizes and larger MAFs lead to a smaller fraction of SNPs estimated to be null. They do however allow for improved quantification of this relationship: For example, we see that the range for 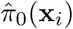 is relatively wide ([0.697,1] for the final smoothed estimate), while the Storey (2002) smoothed estimate of *π*_0_ without covariates is 0.949.

**Figure 2:**
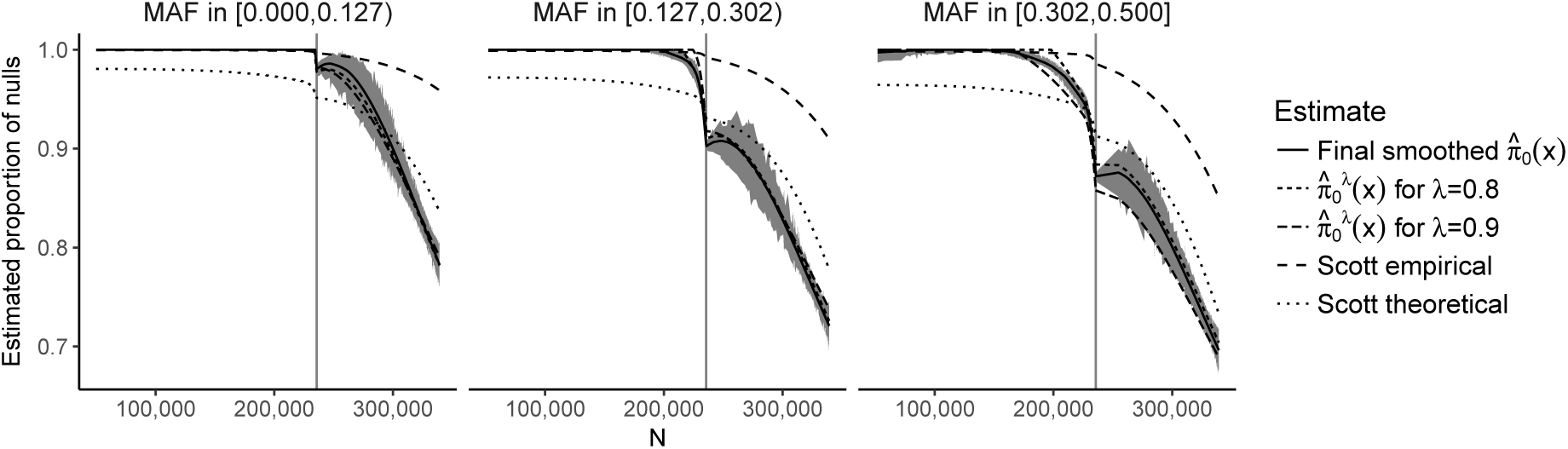
Plot of the estimates of *π*_0_(x_*i*_) against the sample size N, stratified by the MAF categories for a random subset of 50,000 SNPs. The 90% bootstrap intervals for the final smoothed estimates using our approach - based on 100 iterations - are shown in grey. The vertical line represents the median sample size.

The results for the number of SNPs with estimated FDR ≤ 0.05 are given in Table S1. Our approach results in a slightly larger number of discoveries compared to the Storey (2002) and Benjamini and Hochberg (1995) approaches. Due to the plug-in approaches of both our procedure and the one of Storey (2002), all the discoveries from Benjamini and Hochberg (1995) are also present in our approach. The total number of shared discoveries between our method and that of Storey (2002) is 12,740. The Scott et al. (2015) approaches result in either a substantially larger number of discoveries (theoretical null) or a substantially smaller number of discoveries (empirical null). In particular, the number of discoveries for the empirical null is also much smaller than that when using Benjamini and Hochberg (1995). The overlap between the theoretical null and Benjamini and Hochberg (1995) is 12,251; between the theoretical null and our approach it is 13,119.

## 5 Special cases

### 5.1 No covariates

If we do not consider any covariates, the usual estimator 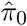 from Eq. (4) can be deduced from applying Algorithm 1 by fitting a linear regression with just an intercept.

### 5.2 Partioning the features

Now assume that the set of features is partitioned into *S* sets, namely that a collection of sets *𝒮* = {*A*_*s*_: 1 ≤ *s* ≤ *S*} is considered such that all sets are non-empty, pairwise disjoint, and have the set of all the features as their union. We consider them ordered for the sake of convenience, for example, the first set in *𝒮* is *A*_1_ et cetera, but note that this ordering does not necessarily have scientific relevance. In the GWAS meta-analysis dataset, the SNPs are partitioned according to their MAFs. Other examples of such partionings include all possible atoms resulting from gene-set annotations or brain regions of interest in a functional imaging analysis, when considering only the genes or voxels that are annotated (Boca et al., 2013). We can consider this in the covariate framework we developed by taking the set *A*_1_ as the “baseline set.” We then consider vectors x_*i*_ of length *S* − 1, with each element corresponding to the remaining sets in *𝒮*, {*A*_2_,…, *A*_*S*_}. We define x_*i*_ = 0 for *i* ∈ *A*_1_ and:

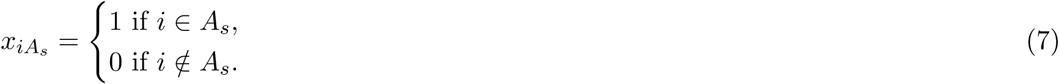

for *i* ∈ *A*_*s*_, *s* ≥ 2. Thus, using the component notation in linear algebra:

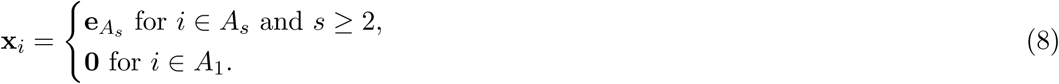

Taking into account the partition, a natural way of estimating *π*_0_(x_*i*_) is to just apply the estimator 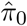 from Eq. (4) to each of the *S* sets:

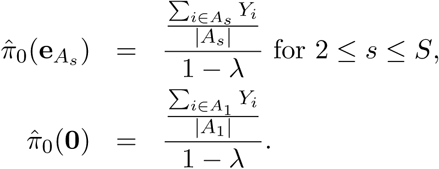

A related idea has been proposed for partitioning hypotheses into sets to improve power (Efron, 2008). These results would be obtained directly from our approach if we considered linear instead of logistic regression and fit a linear regression with an intercept and the covariates x*_i_* in Algorithm 1. As we are considering a logistic regression approach, our results will be slightly different.

## 6 Theoretical results

We now proceed to explore some theoretical properties of the estimator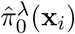. Applying Theorem 3 to Eq. (5), we get that:

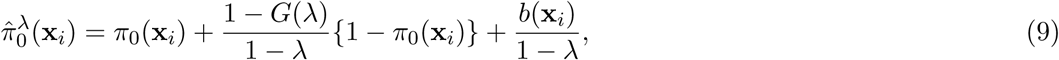

where 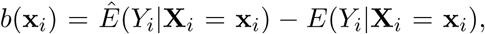, so that *E*{*b*(x_*i*_)} is the bias of 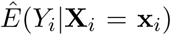 when estimating *E*(Y_*i*_|**X**_*i*_ = x_*i*_). Note that 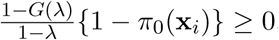, since *λ* ≤ 1,*G*(*λ*) ≤ 1, and *π*_0_(x_*i*_) ≤ 1. Thus, if the bias when estimating *E*(Y_*i*_|**X**_*i*_ = x_*i*_) is positive or negative and small in absolute value, then 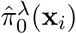 is a conservative estimator of *π*_0_(x_*i*_). For example, if we had considered a correctly specified linear regression model, this would always hold; indeed the case where *π*_0_ is shared by all the features, i.e. in the case of no dependence on covariates, this is shown in Storey (2002). Given that here we are taking 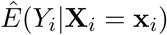 to be the MLE from the logistic regression model, we know that it represents a consistent estimator of *E*(Y_*i*_|**X**_*i*_ = x_*i*_) if the model is correctly specified for *m* → ∞, given certain technical conditions, for instance those specified in Gourieroux and Monfort (1981). Thus, we can show that 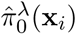 a consistent estimator of 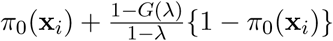 under these same conditions:

### Theorem 5

*Under a correctly specified model and technical regularity conditions*,

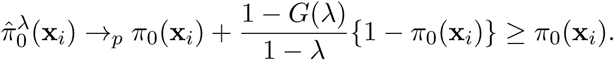

*as m*→∞.

Eq. (9) also leads to 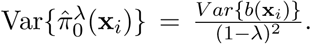. Once again, using the properties of the MLE, under appropriate conditions:

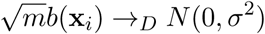

for some *σ*^2^, leading to 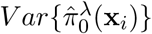 being approximately inversely proportional to m for large values of *m*.

We note that our approach to estimating *π*_0_(x_*i*_) does not place any restrictions on its range. In practice, the values will also be thresholded to be between 0 and 1, as detailed in Algorithm 1. In Result 6, we show that implementing this thresholding decreases the mean squared error of the estimator. The approach is similar to that taken in Theorem 2 in the work of Storey (2002).

**Result 6** *Let*

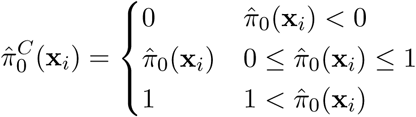

*Then:*

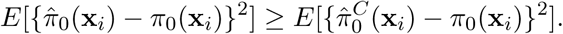

## 7 Simulations

We consider simulations to evaluate the usefulness of our plug-in estimator,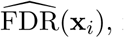, in terms of both controlling the true FDR and having good power - measured by the true positive rate (TPR) - under a variety of scenarios. We consider a nominal FDR value of 5%, meaning that any test with an FDR less than or equal to 5% is considered a discovery. In each simulation, the FDR is calculated as the fraction of truly null discoveries out of the total number of discoveries and the TPR is the fraction of truly alternative discoveries out of the total number of truly alternative features. In the case of no discoveries, the FDR is estimated to be 0.

We consider 4 different possible functions *π*_0_(x_*i*_), shown in Figure 3. Scenario I considers a flat function *π*_0_ = 0.9, to illustrate a case where there is no dependence on covariates and scenarios II-IV are similar to the BMI GWAS meta-analysis. Scenario II is a smooth function of one variable similar to the rightmost panel in Figure 2, scenario III is a function which is smooth in one variable within categories of a second variable - similar to the stratification of SNPs within MAFs - and scenario IV is the same function as in scenario III multiplied by 0.6, to show the effect of having much lower fractions of null hypotheses, respectively higher fractions of alternative hypotheses. The exact functions are given in the Supplementary Materials for this paper. For scenario I we consider fitting a model that is linear in *x*_1_ on the logistic scale, whereas for scenarios II-IV we consider a model that is linear in *x*_1_ and a model that fits cubic splines with 3 degrees of freedom for *x*_1_, both on the logistic scale. For scenarios III and IV, all models also consider different coefficients for the categories of *x*_2_. We set up simulations with independent test statistics for *m* = 1,000 and *m* = 10,000 features and additionally, with dependent test statistics for *m* = 1,000 features and within each setup, different distributions for the alternative test statistics/p-values, the null always assuming a Unif(0,1) distribution. For each combination of factors, we consider 200 simulation runs and obtain the average FDR and TPR over these runs. For each simulation run, we first randomly generated whether each feature was from the null or alternative distribution, so that the null hypothesis was true for the features for which a success was drawn from the Bernoulli distribution with probability *π*_0_(x_*i*_).

**Figure 3:**
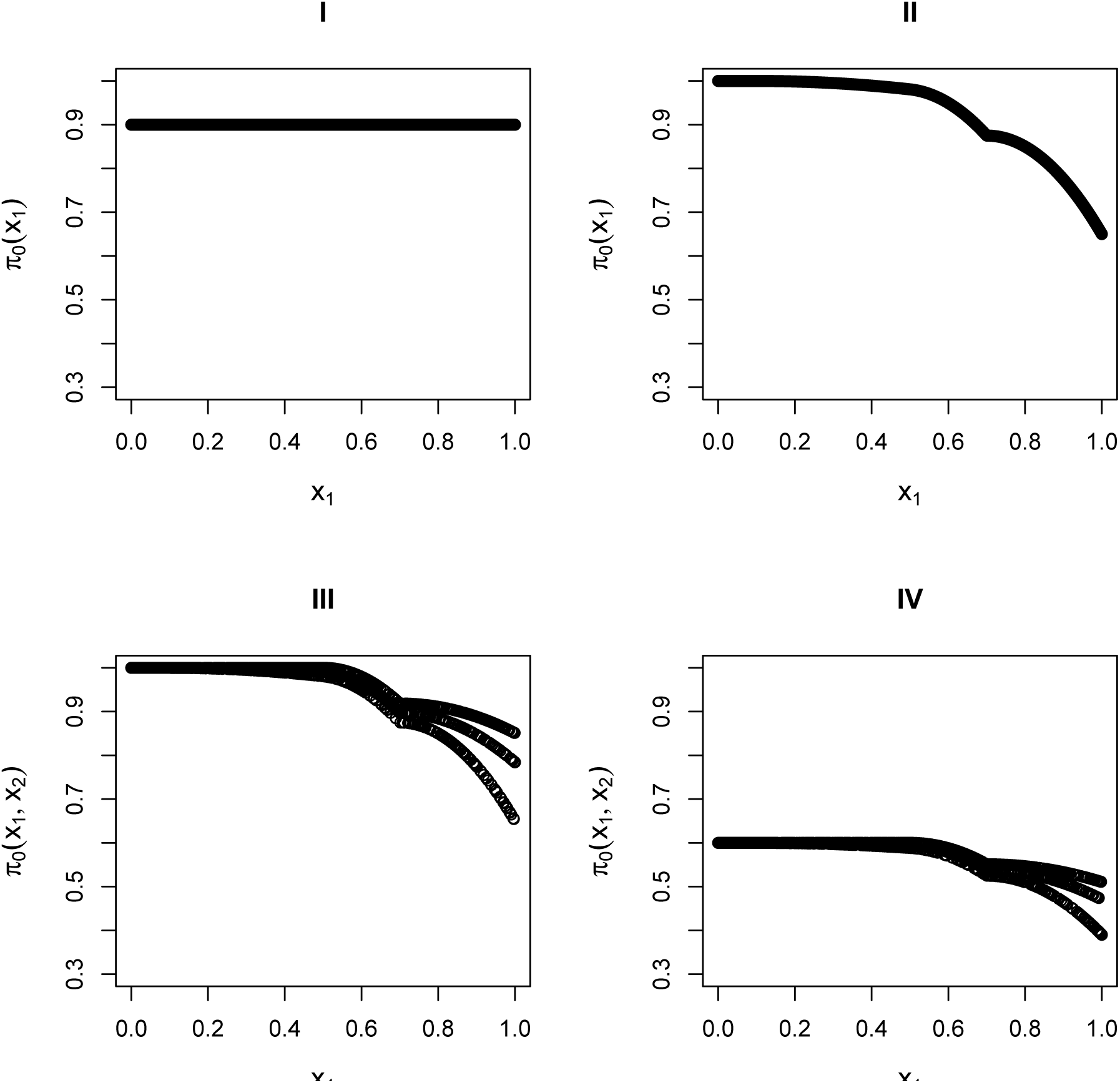
The four simulation scenarios considered for *π*_0_(x_*i*_). Scenarios I and II consider smooth functions of a single covariate, whereas scenarios III and IV consider smooht functions of a single covariate (x_1_) within categories of a second covariate (x

Table 2 and Table 3 consider simulation results for *m* = 1,000 features and *m* = 10,000 features respectively For the Beta(1, 20) simulations, we generated the alternative p-values directly from a Beta(1,20) distribution. For the other simulations, we first generated the test statistics, then calculated the p-values from them. For the normally distributed and t-distributed test statistics, we drew the means *μ*_*i*_ of approximately half the alternative features from a N(*μ* = 3,*σ*^2^ = 1), with the remaining alternative features from a N(*μ* = − 3,*σ*^2^ = 1) distribution, with the mean of the null features being 0. We then drew the actual test statistic for feature *i* from either a N(*μ* = *μ*_*i*_,*σ*^2^ = 1) or T(*μ* = *μ*_*i*_,*df* = 10) distribution (df = degrees of freedom). Note that 10 degrees of freedom for a t-distribution is obtained from a two-sample t-test with 6 samples per group, assuming equal variances in the groups. We also considered chi-squared test statistics with either 1 degree of freedom (corresponding to a test of independence for a 2 × 2 table) or 4 degrees of freedom (corresponding to a test of independence for a 3 × 3 table). In this case, we first drew the non-centrality parameter (ncp_*i*_) from the square of a N(*μ* = 3, *σ*^2^ = 1) distribution for the alternative and took it to be 0 for the null, then generated the test statistics from *χ*^2^(ncp_*i*_ = *μ*_*i*_, *df* = 1 or 4).

**Table 2.** Simulation results for *m* = 1,000 features, 200 runs for each scenario, independent test statistics. “Reg. model” = specific logistic regression model considered, BL = Boca-Leek, Scott T = Scott theoretical null, Scott E = Scott empirical null, BH = Benjamini-Hochberg. A nominal FDR = 5% was considered. Results for the Scott approaches are only presented for scenarios which generate z-statistics or t-statistics.

| $\pi_0(x)$ | Dist. under $H_1$ | Reg. model | FDR % | | | | | TPR % | | | | |
| --- | --- | --- | --- | --- | --- | --- | --- | --- | --- | --- | --- | --- |
|  |  |  | BL | Scott<br>T | Scott<br>E | Storey | BH | BL | Scott<br>T | Scott<br>E | Storey | BH |
| I | Beta(1,20) | Linear | 5.0 |  |  | 5.2 | 3.9 | 0.2 |  |  | 0.2 | 0.1 |
| II | Beta(1,20) | Linear | 4.8 |  |  | 4.8 | 4.1 | 0.2 |  |  | 0.1 | 0.1 |
| II | Beta(1,20) | Spline | 6.5 |  |  | 4.8 | 4.1 | 0.2 |  |  | 0.1 | 0.1 |
| III | Beta(1,20) | Linear | 5.2 |  |  | 5.4 | 5.4 | 0.2 |  |  | 0.2 | 0.2 |
| III | Beta(1,20) | Spline | 6.2 |  |  | 5.4 | 5.4 | 0.3 |  |  | 0.2 | 0.2 |
| IV | Beta(1,20) | Linear | 6.4 |  |  | 5.1 | 3.4 | 12.2 |  |  | 5.4 | 0.3 |
| IV | Beta(1,20) | Spline | 7.9 |  |  | 5.1 | 3.4 | 15.4 |  |  | 5.4 | 0.3 |
| I | Norm | Linear | 5.0 | 5.2 | 6.6 | 4.9 | 4.4 | 51.0 | 50.9 | 49.7 | 50.8 | 49.7 |
| II | Norm | Linear | 5.4 | 5.7 | 8.1 | 5.3 | 4.9 | 48.5 | 63.5 | 61.3 | 47.6 | 47.0 |
| II | Norm | Spline | 5.6 | 5.9 | 8.3 | 5.3 | 4.9 | 49.3 | 63.5 | 61.5 | 47.6 | 47.0 |
| III | Norm | Linear | 5.8 | 5.9 | 9.9 | 5.4 | 5.1 | 45.1 | 60.3 | 57.9 | 44.0 | 43.4 |
| III | Norm | Spline | 5.9 | 6.0 | 10.1 | 5.4 | 5.1 | 45.6 | 60.9 | 58.2 | 44.0 | 43.4 |
| IV | Norm | Linear | 5.0 | 4.9 | 2.4 | 4.7 | 2.8 | 71.6 | 71.8 | 60.6 | 71.2 | 65.4 |
| IV | Norm | Spline | 5.2 | 5.0 | 2.4 | 4.7 | 2.8 | 72.0 | 71.9 | 60.7 | 71.2 | 65.4 |
| I | T | Linear | 5.7 | 21.3 | 23.4 | 5.5 | 4.8 | 15.7 | 55.4 | 56.9 | 15.2 | 13.6 |
| II | T | Linear | 4.8 | 20.7 | 23.8 | 5.0 | 4.4 | 13.0 | 64.5 | 65.5 | 11.6 | 10.6 |
| II | T | Spline | 4.7 | 21.1 | 24.5 | 5.0 | 4.4 | 13.8 | 64.8 | 65.6 | 11.6 | 10.6 |
| III | T | Linear | 6.2 | 26.8 | 31.0 | 5.9 | 5.4 | 9.4 | 54.6 | 54.7 | 8.2 | 7.6 |
| III | T | Spline | 6.8 | 27.3 | 31.3 | 5.9 | 5.4 | 10.0 | 55.2 | 55.3 | 8.2 | 7.6 |
| IV | T | Linear | 5.0 | 9.3 | 2.8 | 4.7 | 2.9 | 52.5 | 72.9 | 44.4 | 52.0 | 40.3 |
| IV | T | Spline | 5.4 | 9.3 | 2.8 | 4.7 | 2.9 | 53.0 | 73.0 | 44.6 | 52.0 | 40.3 |
| I | Chisq 1 df | Linear | 5.0 |  |  | 4.8 | 4.4 | 51.2 |  |  | 50.9 | 49.7 |
| II | Chisq 1 df | Linear | 4.8 |  |  | 4.8 | 4.4 | 48.3 |  |  | 47.1 | 46.3 |
| II | Chisq 1 df | Spline | 5.0 |  |  | 4.8 | 4.4 | 48.9 |  |  | 47.1 | 46.3 |
| III | Chisq 1 df | Linear | 5.0 |  |  | 4.9 | 4.8 | 44.3 |  |  | 43.1 | 42.5 |
| III | Chisq 1 df | Spline | 5.3 |  |  | 4.9 | 4.8 | 44.8 |  |  | 43.1 | 42.5 |
| IV | Chisq 1 df | Linear | 5.1 |  |  | 4.7 | 2.8 | 71.6 |  |  | 71.1 | 65.1 |
| IV | Chisq 1 df | Spline | 5.3 |  |  | 4.7 | 2.8 | 71.9 |  |  | 71.1 | 65.1 |
| I | Chisq 4 df | Linear | 5.3 |  |  | 5.4 | 4.8 | 30.8 |  |  | 30.6 | 29.6 |
| II | Chisq 4 df | Linear | 5.3 |  |  | 5.3 | 5.0 | 28.4 |  |  | 27.5 | 26.7 |
| II | Chisq 4 df | Spline | 5.4 |  |  | 5.3 | 5.0 | 29.2 |  |  | 27.5 | 26.7 |
| III | Chisq 4 df | Linear | 5.9 |  |  | 5.4 | 5.3 | 24.8 |  |  | 24.0 | 23.4 |
| III | Chisq 4 df | Spline | 5.9 |  |  | 5.4 | 5.3 | 25.2 |  |  | 24.0 | 23.4 |
| IV | Chisq 4 df | Linear | 5.1 |  |  | 4.7 | 2.8 | 52.3 |  |  | 51.7 | 44.5 |
| IV | Chisq 4 df | Spline | 5.5 |  |  | 4.7 | 2.8 | 52.7 |  |  | 51.7 | 44.5 |

**Table 3.** Simulation results for *m* = 10, 000 features, 200 runs for each scenario, independent test statistics. “Reg. model” = specific logistic regression model considered, BL = Boca-Leek, Scott T = Scott theoretical null, Scott E = Scott empirical null, BH = Benjamini-Hochberg. A nominal FDR = 5% was considered. Results for the Scott approaches are only presented for scenarios which generate z-statistics or t-statistics.

| $\pi_0(x)$ | Dist. under $H_1$ | Reg. model | FDR % | | | | | TPR % | | | | |
| --- | --- | --- | --- | --- | --- | --- | --- | --- | --- | --- | --- | --- |
|  |  |  | BL | Scott<br>T | Scott<br>E | Storey | BH | BL | Scott<br>T | Scott<br>E | Storey | BH |
| I | Beta(1,20) | Linear | 3.7 |  |  | 3.7 | 3.6 | 0.0 |  |  | 0.0 | 0.0 |
| II | Beta(1,20) | Linear | 3.1 |  |  | 3.1 | 3.0 | 0.0 |  |  | 0.0 | 0.0 |
| II | Beta(1,20) | Spline | 3.1 |  |  | 3.1 | 3.0 | 0.0 |  |  | 0.0 | 0.0 |
| III | Beta(1,20) | Linear | 4.0 |  |  | 3.5 | 3.5 | 0.0 |  |  | 0.0 | 0.0 |
| III | Beta(1,20) | Spline | 4.5 |  |  | 3.5 | 3.5 | 0.0 |  |  | 0.0 | 0.0 |
| IV | Beta(1,20) | Linear | 4.4 |  |  | 4.8 | 2.5 | 1.2 |  |  | 0.5 | 0.0 |
| IV | Beta(1,20) | Spline | 5.0 |  |  | 4.8 | 2.5 | 2.0 |  |  | 0.5 | 0.0 |
| I | Norm | Linear | 5.0 | 5.0 | 5.9 | 5.0 | 4.5 | 50.6 | 50.6 | 52.1 | 50.7 | 49.6 |
| II | Norm | Linear | 4.9 | 5.2 | 5.3 | 4.9 | 4.6 | 48.4 | 63.9 | 62.9 | 47.3 | 46.6 |
| II | Norm | Spline | 4.9 | 5.2 | 5.3 | 4.9 | 4.6 | 48.8 | 64.0 | 63.0 | 47.3 | 46.6 |
| III | Norm | Linear | 4.9 | 5.2 | 5.5 | 4.9 | 4.7 | 44.2 | 60.2 | 59.3 | 43.5 | 43.0 |
| III | Norm | Spline | 4.9 | 5.2 | 5.4 | 4.9 | 4.7 | 44.4 | 60.6 | 59.7 | 43.5 | 43.0 |
| IV | Norm | Linear | 4.8 | 5.0 | 2.3 | 4.8 | 2.8 | 71.3 | 71.8 | 62.2 | 71.2 | 65.3 |
| IV | Norm | Spline | 4.8 | 5.0 | 2.3 | 4.8 | 2.8 | 71.3 | 71.8 | 62.2 | 71.2 | 65.3 |
| I | T | Linear | 5.2 | 21.7 | 20.8 | 5.1 | 4.7 | 14.1 | 55.3 | 53.2 | 14.1 | 12.6 |
| II | T | Linear | 4.6 | 20.0 | 19.9 | 4.9 | 4.5 | 11.5 | 65.7 | 65.4 | 10.2 | 9.2 |
| II | T | Spline | 4.5 | 20.2 | 20.1 | 4.9 | 4.5 | 12.0 | 65.7 | 65.4 | 10.2 | 9.2 |
| III | T | Linear | 4.9 | 24.7 | 26.8 | 5.2 | 5.2 | 6.8 | 62.5 | 63.7 | 6.0 | 5.5 |
| III | T | Spline | 4.8 | 24.8 | 26.9 | 5.2 | 5.2 | 7.0 | 62.6 | 63.9 | 6.0 | 5.5 |
| IV | T | Linear | 4.8 | 9.3 | 1.2 | 4.8 | 2.9 | 51.8 | 72.8 | 28.5 | 51.6 | 40.2 |
| IV | T | Spline | 4.8 | 9.3 | 1.2 | 4.8 | 2.9 | 51.9 | 72.9 | 28.6 | 51.6 | 40.2 |
| I | Chisq 1 df | Linear | 5.0 |  |  | 5.0 | 4.5 | 50.7 |  |  | 50.6 | 49.6 |
| II | Chisq 1 df | Linear | 4.9 |  |  | 5.0 | 4.6 | 48.2 |  |  | 47.2 | 46.4 |
| II | Chisq 1 df | Spline | 4.8 |  |  | 5.0 | 4.6 | 48.6 |  |  | 47.2 | 46.4 |
| III | Chisq 1 df | Linear | 5.0 |  |  | 5.0 | 4.8 | 44.0 |  |  | 43.2 | 42.7 |
| III | Chisq 1 df | Spline | 5.0 |  |  | 5.0 | 4.8 | 44.2 |  |  | 43.2 | 42.7 |
| IV | Chisq 1 df | Linear | 4.8 |  |  | 4.8 | 2.8 | 71.1 |  |  | 71.0 | 65.2 |
| IV | Chisq 1 df | Spline | 4.8 |  |  | 4.8 | 2.8 | 71.2 |  |  | 71.0 | 65.2 |
| I | Chisq 4 df | Linear | 5.0 |  |  | 5.0 | 4.5 | 29.7 |  |  | 29.7 | 28.7 |
| II | Chisq 4 df | Linear | 4.9 |  |  | 5.0 | 4.7 | 28.0 |  |  | 27.1 | 26.5 |
| II | Chisq 4 df | Spline | 4.9 |  |  | 5.0 | 4.7 | 28.4 |  |  | 27.1 | 26.5 |
| III | Chisq 4 df | Linear | 5.2 |  |  | 5.2 | 5.0 | 24.3 |  |  | 23.6 | 23.2 |
| III | Chisq 4 df | Spline | 5.2 |  |  | 5.2 | 5.0 | 24.4 |  |  | 23.6 | 23.2 |
| IV | Chisq 4 df | Linear | 4.7 |  |  | 4.7 | 2.8 | 51.8 |  |  | 51.7 | 44.8 |
| IV | Chisq 4 df | Spline | 4.7 |  |  | 4.7 | 2.8 | 51.9 |  |  | 51.7 | 44.8 |

We compared our approach (BL = Boca-Leek) to the Benjamini and Hochberg (1995) (BH) approach, the Storey (2002) approach as implemented in the `qvalue` package Storey et al. (2015), and both the theoretical and null EB approaches of Scott et al. (2015) (Scott T = theoretical null, Scott E = empirical null), implemented in the `FDRreg` package. The Scott et al. (2015) approaches use z-values, as opposed to the other methods, which use p-values; for the cases of z-statistics and t-statistics, we input these directly into the Scott approaches, while for the remaining simulations we only report results from the remaining three methods.

We see in Tables 2 and 3 that our approach had a true FDR close to the nominal value of 5% in most scenarios. As expected, its performance is better for the larger value of m, with some slight anticonservative behavior for *m* = 1, 000, especially when considering the spline models. The Scott et al. (2015) approaches perform the best in the case where the test statistics are normally distributed, as expected. In particular, the FDR control of the theoretical null approach is also close to the nominal level and the TPR can be 15% higher in absolute terms than that of our approach for scenarios II and III. The empirical null performs less well. However, the Scott et al. (2015) approaches lose control of the FDR when used with t-statistics and are not applicable to the other scenarios. We always see a gain in power for our method over the BH approach, however it is often marginal (1-3%) for scenarios I-III, which have relatively high values of *π*_0_(X_*i*_), which is to be expected, since BH in essence assumes *π*_0_(X_*i*_) = 1. For scenario IV, however, the average TPR may increase by as much as 6% to 11% in absolute terms for *m* = 10,000 while still maintaining the FDR. The gains over the Storey (2002) approach are much more modest, as expected (0-2% in absolute terms while maintaining the FDR for *m* = 10,000). We also compare the empirical means of the estimates of *π*_0_(X_*i*_) over the 200 simulation runs compared to the true values of *π*_0_(X_*i*_) for the normally-distributed and t-distributed independent test statistics in Figures S1 - S4. We note that for the t-distributed statistics, the Scott theoretical null estimate is less conservative than ours in scenario I (we considered only the theoretical, not the empirical null for the Scott approach in the plots, given that the theoretical null had much better properties in our simulations, as seen from Tables 2 and 3). For scenarios II and III, the Scott theoretical null was more anti-conservative for lower values of *π*_0_(X_*i*_), leading to much higher FDRs in Tables 2 and 3.

Tables S2 and S3 consider simulation results for *m* = 1,000 features and several dependence structures for the test statistics. We considered multivariate normal and t distributions, with the means drawn as before and block-diagonal variance-covariance matrices with the diagonal entries equal to 1 and a number of blocks equal to either 20 (50 features per block) or 10 (100 features per block). The within-block correlations, *ρ*, were set to 0.2, 0.5, or 0.9. As expected, the FDR was generally closer to the nominal value of 5% for 20 blocks than for 10 blocks, as 20 blocks leads to less correlation. Increasing *ρ* also leads to worse control of the FDR. These same trends are also present for the Scott et al. (2015) approaches, but generally with worse control. Furthermore, for *ρ* = 0.5, the empirical null leads to errors in 1% or fewer of the simulation runs; however, for *ρ* = 0.9 it leads to errors in as many as 33% of the runs. In contrast, Storey (2002) shows estimated FDR values closer to 5% and results in a single error for *ρ*= 0.9 and 10 blocks for the t distribution. We also note that the TPR is generally very low for the multivariate t distributions, except in scenario IV.

## 8 Reproducibility

All analyses and simulations in this paper are fully reproducible and the code is available on Github at: https://github.com/SiminaB/Fdr-regression

## 9 Discussion

Here we have introduced an approach to estimating false discovery rates conditional on covariates in a multiple testing framework, by first estimating the proportion of true null hypotheses via a regression model and then using this in a plug-in estimator. Our motivating case study considers a GWAS meta-analysis of BMI-SNP associations, where we are interested in adjusting for sample sizes and allele frequencies of the individual SNPs. Using extensive simulations, we compared our approach to FDR regression as proposed by Scott et al. (2015), as well as to the approaches of Benjamini and Hochberg (1995) and Storey (2002), which estimate the FDR without covariates. While the Scott et al. (2015) approaches outperform our approach for normally-distributed test statistics, which is one of modeling assumptions therein, that approach tends to lose FDR control for test statistics from the t-distribution and cannot be applied in cases where the test statistics come from other distributions, such as the chi-squared distribution, which may arise from commonly performed analyses. In general, our method provides the flexibility of performing the modeling at the level of the p-values. Our approach always shows a gain in true positive rate over Benjamini and Hochberg (1995), which is often limited, but was as high as 6%-11% in our simulations for low values of the prior probabilities. While the gains over the Storey (2002) approach were more modest, our method allows for improved flexibility in modeling, as evidenced in Figures S1 - S4. It may also be the case that estimating the proportion of true null hypotheses as a function of covariates is of interest. We further show that control of the FDR is maintained in the presence of moderate correlation between the test statistics.

Applying our estimator to GWAS data from the GIANT consortium demonstrated that, as expected, the estimate of the fraction of null hypotheses decreases with both sample size and minor allele frequency. It is a well-known and problematic phenomenon that p-values for all features decrease as the sample size increases. This is because the null is rarely precisely true for any given feature. One interesting consequence of our estimates is that we can calibrate what fraction of p-values appear to be drawn from the non-null distribution as a function of sample size, potentially allowing us to quantify the effect of the “large sample size means small p-values” problem directly. Using an FDR cutoff of 5%, our approach leads to 13,384 discoveries, compared to 12,771 from the Storey (2002) method; given the fact that they are both multiplicative factors to the Benjamini and Hochberg (1995) approach, which in effect assumes the proportion of true null hypotheses to be 1, they both include the 12,500 discoveries using this approach. Thus, our approach leads to additional insights due to incorporating modeling of the fraction of null hypotheses on covariates, as well as to a number of new discoveries. By contrast, the Scott et al. (2015) approach leads to very different results based on whether the theoretical null or empirical null is assumed.

We note that our approach relies on a series of assumptions, such as independence of p-values and independence of the p-values and the covariates conditional on the null or alternative. Assuming that the p-values are independent of the covariates conditional on the null is also an assumption used in Ignatiadis et al. (2016). Therein, diagnostic approaches for checking this assumption are provided, namely examining the histograms of p-values stratified on the covariates. In particular, it is necessary for the distribution to be approximately uniform for larger p-values. We perform this diagnostic check in Figure S5 and note that it appears to hold approximately. The slight conservative behavior seen for smaller values of N in Figures 1 and S5 may be the result of publication bias, where studies with borderline significant p-values become part of larger meta-analyses. It is interesting that the estimated proportion of nulls in Figure 2 also starts decreasing substantially right at the median sample size (of 235,717). This may also be due to the same publication bias.

In conclusion, our approach shows good performance across a range of scenarios and allows for improved interpretability compared to the Storey (2002) method. In contrast to the Scott et al. (2015) approaches, it is applicable outside of the case of normally distributed test statistics. It always leads to an improvement in estimating the true positive rate compared to the now-classical Benjamini and Hochberg (1995) method, which becomes more substantial when the proportion of null hypotheses is lower. While in very high correlation cases, our method does not appropriately control the FDR, we note that in practice methods are often used to account for such issues at the initial modeling stage, meaning that we generally expect good operating characteristics for our approach. In particular, for GWAS, correlations between sets of SNPs (known as linkage disequilibrium) are generally short-range, being due to genetic recombination during meiosis (Frazer et al., 2007); longer-range correlations can result from population structure, which can be accounted for with approaches such as the genomic control correction (Devlin and Roeder, 1999) or principal components analysis (Price et al., 2006). For gene expression studies, batch effects often account for between-gene correlations; many methods exist for removing these, including Johnson et al. (2007); Leek and Storey (2007) and Leek (2014). We also note the subtle issue that the accuracy of the estimation is based on the number of features/tests considered, not on the sample sizes within the tests. Thus, our “large-sample” theoretical results are to be interpreted within this framework. In our simulations, for example, we see that using 10,000 rather than 1,000 features improved the FDR control. We note that our motivating data analysis had over 2.5 million features and that many high-dimensional problems have features in the tens of thousands or higher. A range of other applications for our methodology are also possible by adapting our regression framework, including estimating false discovery rates for gene sets (Boca et al., 2013), estimating science-wise false discovery rates (Jager and Leek, 2013), or improving power in high-throughput biological studies (Ignatiadis et al., 2016). Thus, this is a general problem and as more applications accumulate, we anticipate our approach being increasingly used to provide additional discoveries and scientific insights.

